# Microwave emission from honeybees

**DOI:** 10.1101/2025.03.24.644736

**Authors:** Igor Goryanin, Sergey Vesnin, Xavier Lair, Irina goryanin

## Abstract

Honeybees exhibit complex behaviours in foraging, navigation, and hive maintenance, all regulated by circadian rhythms and influenced by environmental factors, including electromagnetic fields. This paper investigates the role of passive microwave radiometry (MWR) and IR in studying honeybee behaviour, particularly about their internal temperature regulation and circadian rhythms. Experimental data demonstrated that honeybees maintain a stable internal temperature despite environmental fluctuations, with microwave emissions showing a greater temperature amplitude than infrared measurements, and unexpected oscillation in the low Hz range. The experiments confirmed the presence of circadian rhythms in bees under controlled light conditions, and 12-hour oscillations with no lights, contributing to the understanding of their thermoregulatory processes. Additionally, the role of cryptochromes in bee navigation and magnetoreception, potentially mediated by quantum phenomena, is discussed. This interdisciplinary approach also explores the potential practical applications of bee-inspired technologies, including MWR-based health monitoring systems for bee colonies, and advances in quantum computing models.

## Introduction

Honeybees *(Apis mellifera)* play a critical role in ecosystems as pollinators, supporting the reproduction of countless plant species and sustaining global biodiversity. Their behaviour, characterized by intricate foraging patterns, social communication, and precise navigation, has been the focus of extensive biological research. Recent advancements in molecular biology, neuroethology, and environmental sensing technologies have shed light on the underlying mechanisms that govern these behaviours. Central to their daily routines are circadian rhythms, which regulate various aspects of bee activity, including foraging, hive maintenance, and brood care [1,2]

Circadian rhythms in honeybees are closely linked to environmental light cycles and temperature changes, with cryptochromes—light-sensitive proteins—playing a crucial role in synchronizing internal biological clocks [3]. These rhythms not only influence behavioural patterns but also aid in bees’ remarkable navigation abilities, enabling them to orient themselves using the sun, polarized light, and possibly Earth’s magnetic field [4,5]. The latter has drawn attention to the potential role of quantum phenomena in magnetoreception, where cryptochromes may act through quantum coherence and entanglement, providing bees with a biological compass[6].

In parallel with advances in understanding bee physiology and behaviour, there has been growing interest in the application of microwave radiometry (MWR) as a non-invasive tool for studying bees. MWR can measure the internal temperatures of bees, offering insights into their thermoregulation and physiological responses to environmental changes[2]. Previous studies have demonstrated the potential of IR for measuring circadian temperature rhythms. These experiments, conducted under controlled conditions, provide valuable data on how bees maintain a stable internal temperature despite fluctuations in ambient temperatures, reinforcing the importance of circadian regulation in their daily cycles [7,8]

This paper aims to explore the intersection of honeybee behaviour, circadian rhythms, and [9], focusing on the potential role of quantum biological mechanisms in bee navigation.

## Materials and methods

So far, everyone is convinced that all the heated bodies, and really all the bodies that surround us, emit electromagnetic waves in the microwave range. So, everything around us emits a noise signal where power is proportional to the temperature. It is assumed, that In the microwave range, the power of thermal radiation is linearly correlated to body temperature (Eq. 1).

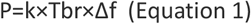

where P is noise power at the output antenna; Tbr is brightness temperature; k is the Boltzmann constant; and Δf is receiver bandwidth. The received power is about 10^-12^ at a median frequency of 4.0Ghz and Δf =600Mhz.

However, recent studies show that, in many cases, non-thermal radiation of living systems in the microwave range also takes place. To better understand the nature of the phenomena several *in vitro* experiments on cell lines [12], enzymes [13,14] and proteins [15-17] were performed using RTMM device. Commercially available equipment:MMWR2020 (former RTM-01-RES) (http://www.mmwr.co.uk) (Fig. 1), has been developed for early diagnosis of disease [18-26]. We decided to use this device to monitor MWR and IR emissions from bees and compare them.

**Figure 1.**
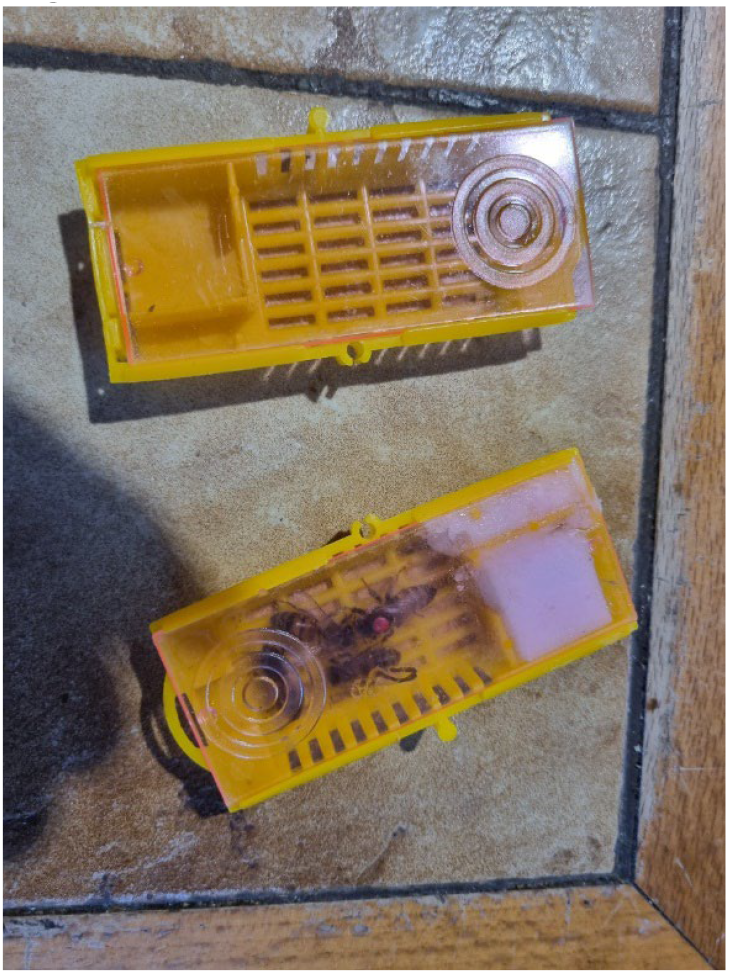
Boxes. Empty and with honeybees and sugar

Initially, we run experiments using boxes with bees. Each box contains 10 bees (Fig. 1). The boxes were attached to the sensor (Fig. 2) and wrapped by magnetic shield tissue to avoid interference (Fig.3)

**Figure 2.**
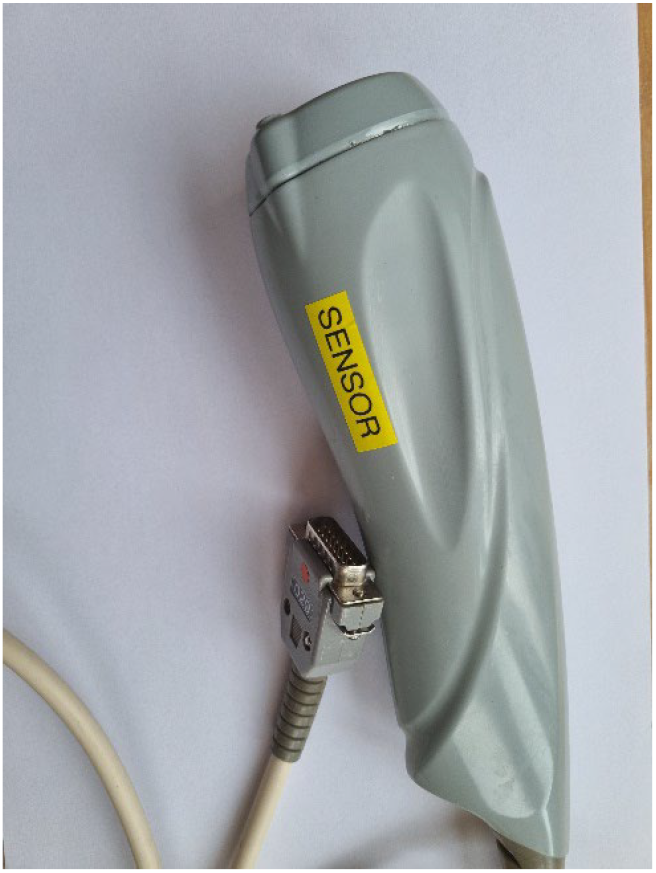
Combined Microwave/IT sensor

**Figure 3.**
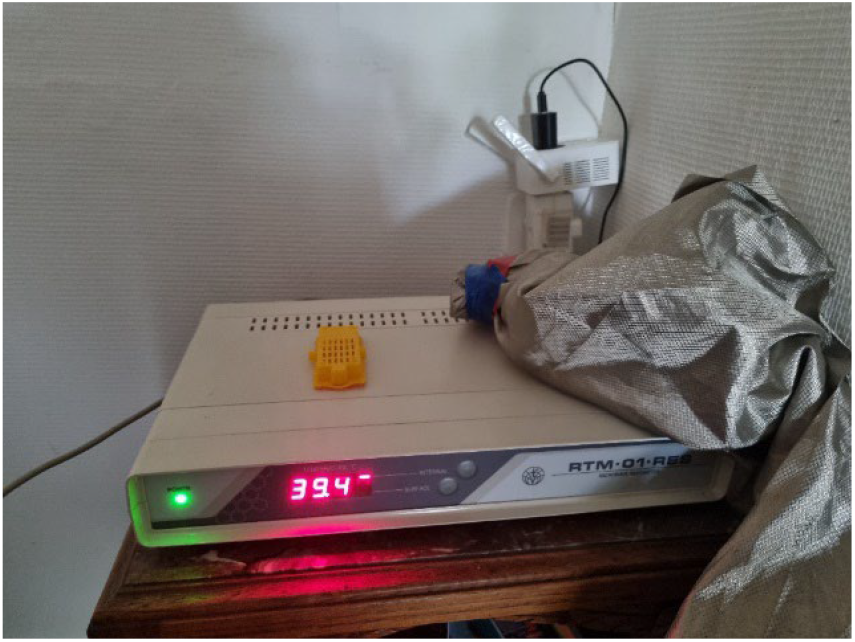
The experimental setup. Digital Unit and sensor with boxes wrapped

Next, we ran experiments using the same device but inserted it into the hive in the garden (Fig. 4) and monitored 24 hours. The colony chosen for this experiment comprises about 4-5 frames of bees, in a Dadant hive, or approx. 10.000 bees. It is a colony with a young queen at the beginning of laying, resulting from a division of a wintered colony. These bees of “Buckfast” origin were chosen because they are not very aggressive. A more developed colony in honey production would pose other difficulties. All experiments were performed 3 times.

**Figure 4.**
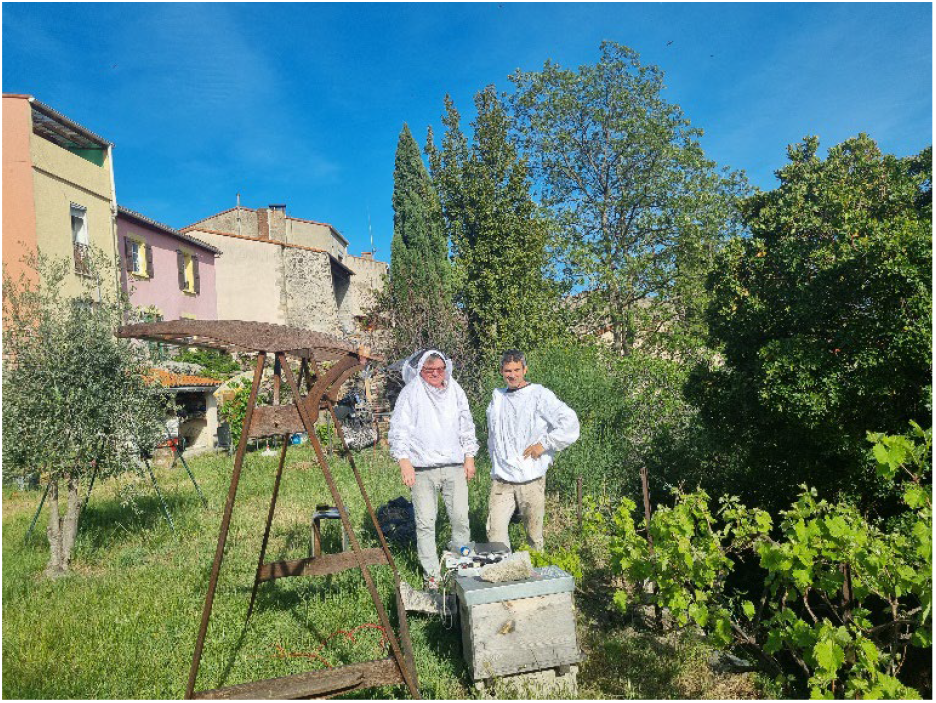
Experimental setup in the garden. There are co-authors (Prof Igor Goryanin, Dr Xavier Lair^)^ on this figure. May 2023

## Results

Initially, control experiments were conducted using three empty boxes (Fig. 5). There is no IR and MWR and IR emissions are stable with no fluctuations.

**Figure 5.**
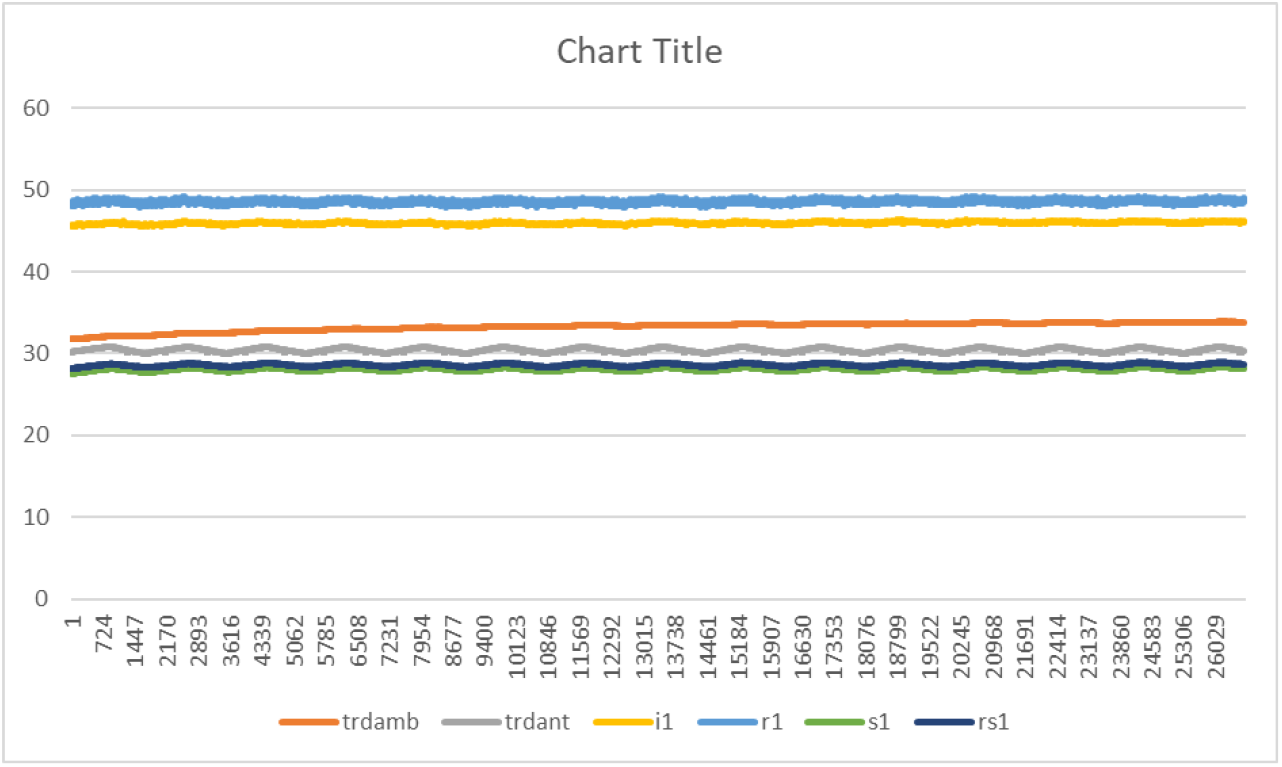
Control Experiment. Six empty boxes were covered by electromagnetic field protection tissue. Y-axes (°C) Trdamb - ambient temperature, trant – device antenna temperature, i1 - compensated microwave emissions, r1 -raw microwave emissions, rs1 - raw IR emissions, s1 - compensated IR emissions .

X – axes. Point number (points correspondent to every 0.25 sec measurements*). 724 corresponds to 724/4=181 sec*

Fig 6 show rhythms in 6 boxes The period of oscillations is about 12 hours.

**Figure 6.**
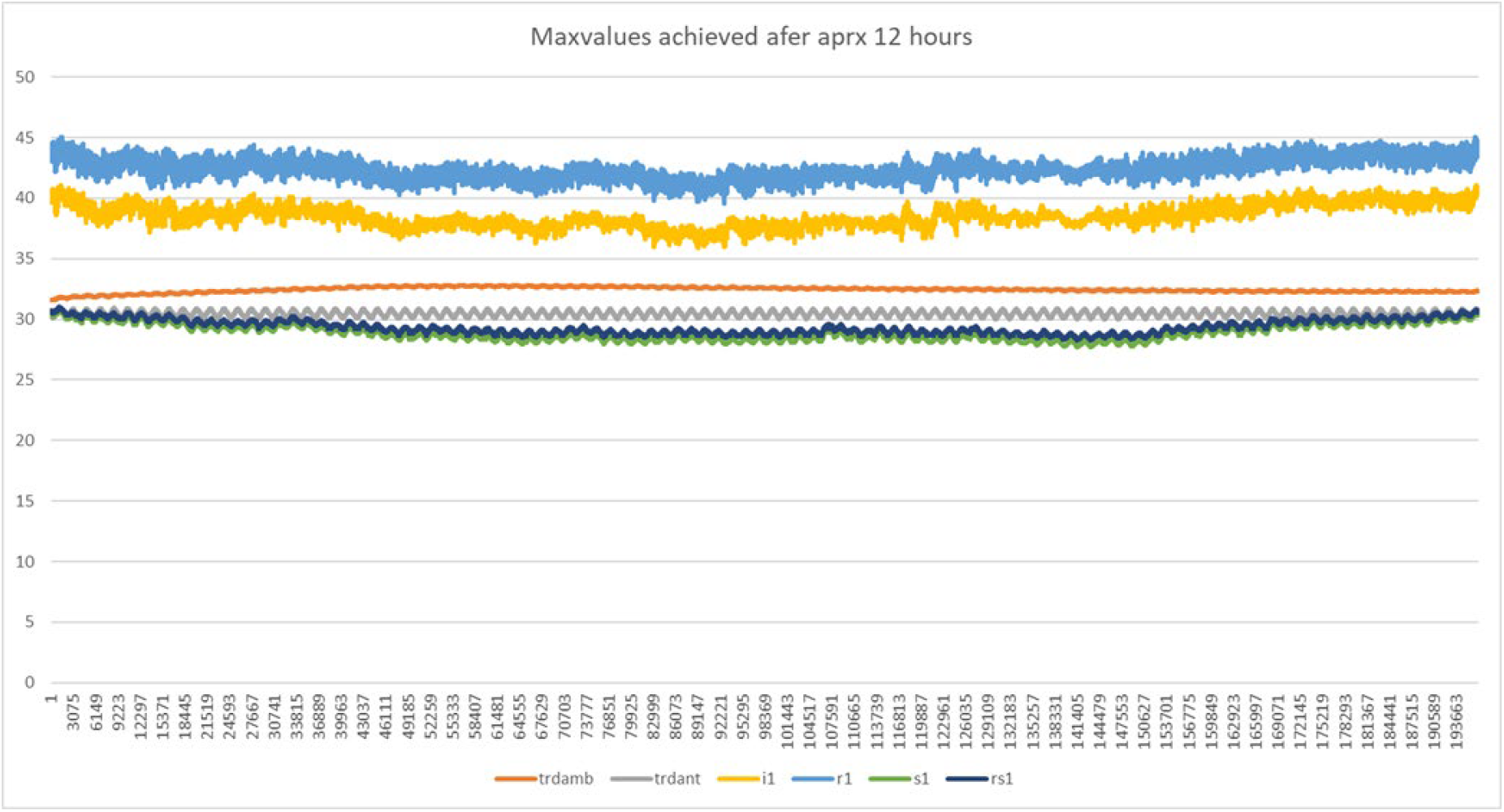
12 hours oscillations in covered 6 boxes with bees and sugar, and no light

Fig 6 shows emissions from 6 boxes with bees. Boxes covered by non-transparent light electromagnetic shield tissue. Measurement every 250 msec. Microwave emission shows oscillations with average, 33°C frequency 1-5Hz, and amplitude 2°C, IR emission show lower temperature, average 30.5°C, and no law frequency oscillations. Twelve hours rhythm were observed in both in MWR and IR ranges with IR at max 31°C, min 29°C. MWR min 39°C and max 45°C.

Fig 7 shows 12 hours oscillations of microwave emissions 33°max at 2am and 2pm, min 28°C are 8am.

**Figure 7.**
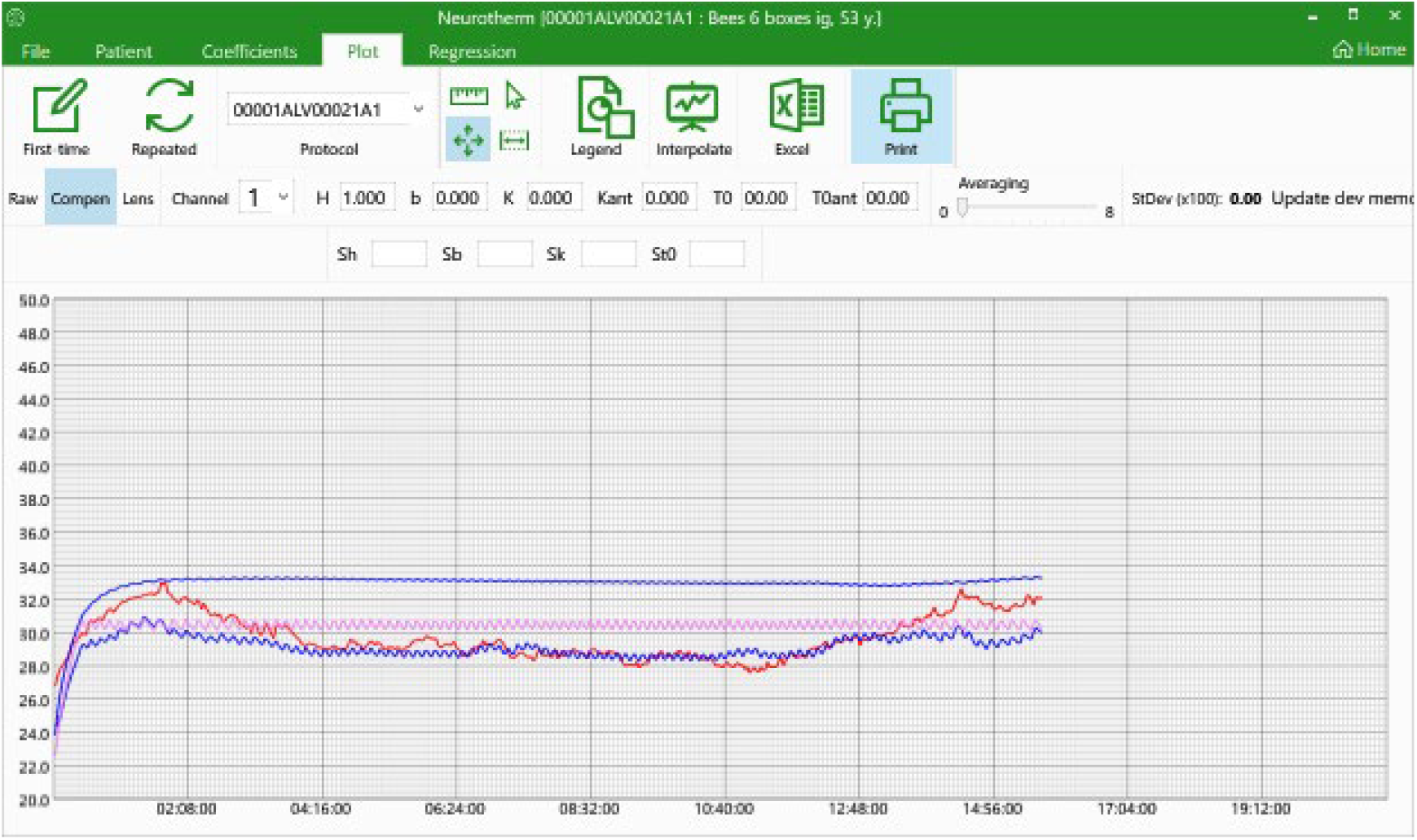
Controlling Software front-en to setup and monitor experiments. Six boxes of bees.

Y-axes: Blue -device temperature, Pink - sensor temperature. Red – MWR emissions, dark blue – IR emission X -axes: time (hours).

Fig. 8 Shows experiments where IR/MWR emissions were measured inside hive during 24 hours. Circadian rhythm is observed within 24 hours period. There were 12 hour rhythms. Low frequency oscillation for both IR/MWR emissions. A large difference in amplitude and absolute values were observe between IR and MWR emissions. Much high emissions in MWR range.

**Figure 8.**
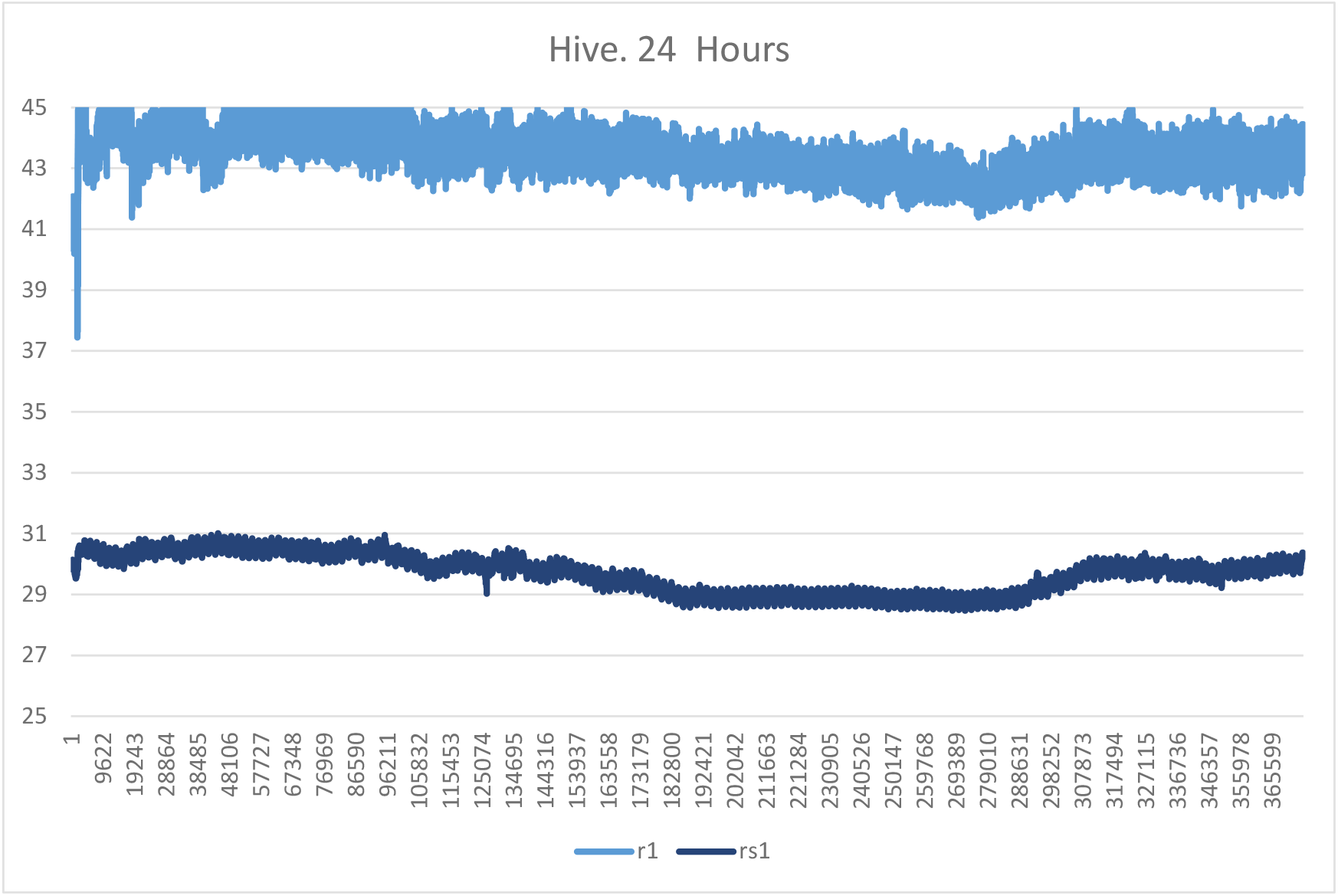
Real Hive Experiments. Circadian Rhythms in honeybees. The difference in amplitude MWR emissions >4°C, and in IR 2°C degrees. Light blue – MWR, Dark blue - IR

## Discussion

Honeybees exhibit a range of circadian behaviours, such as foraging, hive maintenance, and sleep, which can be used to monitor their circadian rhythms. Bees are known to maintain a constant body temperature within a narrow range. Temperature changes can affect the timing and intensity of certain behaviours, such as foraging and brood rearing, which are regulated by the circadian clock.

One way to measure temperature in honeybees is by using thermocouples or thermistors, which are small sensors that can be attached to the bee’s body or placed in the hive. These sensors can record temperature changes over time, allowing researchers to analyse the circadian temperature rhythm of the bees.

In addition to direct temperature measurements, researchers can also use infrared cameras or thermal imaging to monitor hive temperature and activity patterns. This non-invasive technique can provide a detailed view of how temperature changes affect the behaviour of individual bees and the hive. There are many publications on the topic of honeybee circadian rhythms that use temperature measurements and other techniques [21,22]

Our findingd highlight the significant role of microwave radiometry (MWR) in studying honeybee physiology and behaviour, particularly in relation to circadian rhythms and temperature regulation. The MWR experiments demonstrated that honeybees maintain a stable internal temperature despite fluctuations in ambient conditions, with the bees showing a greater amplitude of emission variation when measured via microwaves compared to infrared. This finding aligns with prior research on the thermoregulation abilities of honeybees, which suggests that bees are capable of precisely regulating their body temperature within a narrow range to optimize foraging efficiency and hive maintenance [2].

The MWR results in this paper complement these findings by providing a novel method for monitoring circadian temperature fluctuations, supporting the hypothesis that bees express robust circadian rhythms even in the absence of light. This observation is crucial, as it suggests that temperature, rather than light alone, could be an important environmental Zeitgeber (time-giver) for bees [23-25].

The role of cryptochromes in honeybee circadian regulation and magnetoreception has been a growing area of interest in recent years. Cryptochromes are light-sensitive proteins that play a central role in maintaining circadian rhythms in many organisms, including bees, and are thought to mediate responses to the Earth’s magnetic field, which is essential for their navigation. The involvement of cryptochromes in magnetoreception is potentially linked to quantum coherence and the radical pair mechanism, as shown in studies of migratory birds. Although direct evidence for quantum effects in bees is still under investigation, the MWR experiments could open the door for further exploration into how microwave fields might affect or reveal the quantum biological processes in honeybee navigation. [26-29]

Additionally, the potential impact of electromagnetic pollution on bee navigation and behaviour must be considered. With increasing exposure to anthropogenic electromagnetic radiation from mobile devices, Wi-Fi, and other sources, there is growing concern that this environmental noise could disrupt the cryptochrome-based magnetoreception system in bees [30]. Studies have suggested that exposure to electromagnetic fields can interfere with the radical pair mechanism, potentially leading to navigation errors and contributing to colony collapse disorder (CCD) [31]. The MWR results provide a non-invasive method for further investigating these effects, offering a valuable tool for understanding how environmental electromagnetic fields may influence bee physiology and behaviour.

Bees and cryptochromes have an interesting relationship, particularly in the context of navigation and possibly magnetoreception. Like migratory birds, bees are thought to use cryptochromes to sense magnetic fields, which may help them navigate over large distances to locate flowers and return to their hives. Additionally, cryptochromes are involved in regulating circadian rhythms in bees, influencing their daily activity patterns, foraging behaviour, and hive maintenance.

Bees, like many other animals, possess an extraordinary ability to navigate over large distances. They use a combination of the sun, polarized light, visual landmarks, and, potentially, Earth’s magnetic field to orient themselves. Cryptochromes play a crucial role in this navigation ability, particularly through magnetoreception. Cryptochromes in bees are thought to be sensitive to Earth’s magnetic field through a mechanism known as the radical pair mechanism. When cryptochromes absorb blue light, they undergo a chemical reaction that produces radical pairs (molecules with unpaired electrons). The behavior of these radicals, specifically their spin states, can be influenced by Earth’s magnetic field, providing directional information to the bee.

The role of cryptochromes in bee navigation potentially places bees within the field of quantum biology. As with birds, the radical pair mechanism in cryptochromes could involve quantum coherence, meaning that quantum mechanical effects, such as entanglement, may be present in biological processes.

Quantum effects might allow cryptochromes to remain sensitive to weak magnetic fields over time, helping bees detect small variations in Earth’s magnetic field for navigation. This would make bees one of the examples where quantum mechanics might directly influence animal behavior, although more research is needed to confirm the extent of this effect.

Bees are capable of producing oscillations in the 1-5 Hz range for a variety of reasons related to their communication, movement, and behavior. Here are a few possible explanations:

Bees’ wing beats can produce oscillatory frequencies when they are in flight or hovering. While their wingbeat frequency is much higher (around 200-230 Hz for honeybees), the movements of their bodies or thorax might produce lower frequency oscillations in the 1-5 Hz range during different flight modes or behaviors, such as slow maneuvering or specific social interactions.

Bees use vibration signals to communicate within the hive. Some of these signals, like the “tremble dance” or “shaking signals,” involve periodic body movements that could generate oscillations within the 1-5 Hz range. These vibrational signals can be transmitted through the comb, enabling communication between bees in a dark, crowded environment.

When bees perform collective activities (e.g., fanning their wings to ventilate the hive or creating vibrations for communication), these collective motions could lead to oscillations in the hive structure itself. Resonance effects from many bees working in sync might create low-frequency vibrations in this range.

Bees can engage in thermoregulation by clustering together and vibrating their thoraxes. This group behavior might produce low-frequency oscillations that could fall within the 1-5 Hz range as they maintain hive temperature, especially during cold periods.

Dual circadian rhythms, which we found, where an organism exhibits two distinct peaks of activity in a 24-hour cycle, have been observed in various animals and insects. This phenomenon often involves **two oscillators**: one associated with **morning activity** and another with **evening activity**, sometimes reflecting adaptation to environmental or social cues. They were found in Fruit Flies (*Drosophila melanogaster*), Crickets (*Teleogryllus commodus*), Bees (*Apis mellifera*), Ants (*Temnothorax rugatulus*), Moths (*Spodoptera littorali*s) [32-36].

## Conclusion

We have found that the 24 rhythms occur in a hive which is connected to day and night and linked to the pollinated plants.

Strikingly, bees in the box covered by non-transparent electromagnetic shield tissue with constant access to sugar exhibit 12-hour oscillations in microwave emissions. While IR and MWR emissions have the same period relative amplitude of oscillation in MWR was significantly higher. It could be explained by the fact that the origin of circadian oscillations is MWR emission and IR emissions have a secondary role (like in microwave ovens). In addition, there were Hz oscillations in the Microwave range and but not such oscillations in IR which confirms our hypothesis.

The results of this study emphasize the potential of MWR as an effective non-invasive tool for understanding honeybee physiology, particularly in relation to their thermoregulation and circadian rhythms. MWR has shown to be a valuable method for tracking the internal temperature fluctuations in bees, confirming that bees maintain a stable temperature despite environmental changes. This stability is closely tied to their circadian rhythms, which govern essential behaviours like foraging and hive maintenance

By utilizing MWR to track hive temperature and health, early signs of colony stress, disease, or infestations can be detected. This technology could also be expanded to study the effects of climate change and habitat degradation on bee populations, ensuring their vital role in pollination and ecosystem stability remains protected [37].

## Supporting information

24 hours oscillations

## References

1. Moore, D., & Rankin, M. A. (1985). Circadian locomotor rhythms in individual honeybees. Physiological Entomology, 10(2), 191–197. 10.1111/j.1365-3032.1985.tb00034.x.

2. Stabentheiner, A., Kovac, H., & Brodschneider, R. (2010). Honeybee colony thermoregulation: Brood nest temperature regulation in response to changing environmental conditions. PLoS ONE, 5(1), e08324. 10.1371/journal.pone.0008324.

3. Emery, P., So, W. V., Kaneko, M., Hall, J. C., & Rosbash, M. (1998). Cry, a Drosophila clock and light-regulated cryptochrome, is a major contributor to circadian rhythm resetting and photosensitivity. Cell, 95(5), 669–679. 10.1016/S0092-8674(00)81638-4.

4. Dyer FC. The biology of the dance language. Annu Rev Entomol. 2002;47:917–49. doi: 10.1146/annurev.ento.47.091201.145306. PMID: 11729095.

5. Menzel R, Greggers U. The memory structure of navigation in honeybees. J Comp Physiol A Neuroethol Sens Neural Behav Physiol. 2015 Jun;201(6):547–61. doi: 10.1007/s00359-015-0987-6. Epub 2015 Feb 24. PMID: 25707351.

6. Ritz T, Thalau P, Phillips JB, Wiltschko R, Wiltschko W. Resonance effects indicate a radical-pair mechanism for avian magnetic compass. Nature. 2004 May 13;429(6988):177–80. doi: 10.1038/nature02534. PMID: 15141211.

7. Przemysław Grodzicki, Michał Caputa, Photoperiod influences endogenous rhythm of ambient temperature selection by the honeybee Apis mellifera, Journal of Thermal Biology, Volume 37, Issue 8, 2012, Pages 587–594,ISSN 0306-4565, 10.1016/j.jtherbio.2012.07.005.

8. Eban-Rothschild, A. D., Bloch, G. (2012) Circadian Rhythms and Sleep in Honey Bees, chapter 1.3., p. 31–46 in “ Honeybee Neurobiology and Behavior”,

9. Goryanin, I., Vesnin, S., Lair, X., & Goryanin, I. (2023). Microwave radiometry as a tool for monitoring circadian rhythms in honeybees. Journal of Insect Physiology, (In press).

10. D. Ivanov, et al. Monitoring of brightness temperature of suspension of follicular thyroid carcinoma cells in SHF range by radiothermometry Patol. Fiziol. Eksp. Ter., 60 (2016), pp. 174–177

11. Y.D. Ivanov, et al. Monitoring of microwave emission of HRP system during the enzyme functioning Biochem. Biophys. Rep., 7 (2016), pp. 20–25

12. Y.D. Ivanov, et al. The registration of a biomaser-like effect in an enzyme system with an RTM sensor J. Sensors (2019), 10.1155/2019/7608512

13. 13 I. Goryanin, et al. Monitoring protein denaturation using passive microwave radiometry

14. 14 Ivanov, Yu.D., Malsagova K.A., Tatur, V.Yu., Vesnin, S.G., Ivanova, N.D., Ziborov, V.S. SHF radiation from albumin solution upon external excitation, Patologicheskaia fiziologiia i eksperimental’naia terapiia (Pathological Physiology and Experimental Therapy, in Russian). 60 (3), 101–104 (2016).

15. 15 N.A. Livanos, et al. Design and interdisciplinary simulations of a hand-held device for internal-body temperature sensing using microwave radiometry IEEE Sensors J., 18 (2018), pp. 2421–2433

16. 16 S. Karabetsos, et al. Development of the RF front-end of a multi-channel microwave radiometer for internal body temperature measurements J. Phys. Conf. Ser., 637 (2015), Article 012010

17. 17 R.S. Akki, et al. Multi-physics modeling to study the influence of tissue compression and cold stress on enhancing breast tumor detection using microwave radiometry Bioelectromagnetics, 40 (2019), pp. 260–277

18. 18 M. Sedankin, et al. Development of a miniature microwave radiothermograph for monitoring the internal brain temperature Eastern Eur. J. Enterprise Technol, 3 (2018), pp. 26–36

19. 19 A.G. Gudkov, et al. Use of multichannel microwave radiometry for functional diagnostics of the brain Biomed. Eng., 53 (2019), pp. 108–111

20. 20 M.K. Sedankin, et al. Antenna applicators for medical microwave radiometers Biomed. Eng., 52 (2018), pp. 235–238

21. 21 Giannoni-Guzmán MA, Perez Claudio E, Aleman-Rios J, Diaz Hernandez G, Perez Torres M, Melendez Moreno A, Loubriel D, Moore D, Giray T, Agosto-Rivera JL. The role of temperature on the development of circadian rhythms in honey bee workers. PeerJ. 2024 Mar 15;12:e17086. doi: 10.7717/peerj.17086. PMID: 38500530; PMCID: PMC10946391.

22. Silvio P. Oliveira et al. “Circadian regulation in the honeybee brain is revealed by mutual information analysis” by in Communications Biology (2019).

23. Southwick, E. E. (1982). Metabolic energy of intact honey bee colonies. Comparative Biochemistry and Physiology Part A: Physiology, 72(1), 399–405. 10.1016/0300-9629(82)90344-4.

24. Moore, D., & Rankin, M. A. (1985). Circadian locomotor rhythms in individual honeybees. Physiological Entomology, 10(2), 191–197. 10.1111/j.1365-3032.1985.tb00034.x.

25. Bloch, G. (2010). The social clock of the honeybee. Journal of Biological Rhythms, 25(4), 307–317. 10.1177/0748730410379406.

26. Fedele, G., Green, E. W., Rosato, E., & Kyriacou, C. P. (2014). Magnetic field effects on cryptochrome-dependent responses in Drosophila. Nature Communications, 5, 4391. 10.1038/ncomms5391.

27. Menzel, R., & Greggers, U. (2015). The memory structure of navigation in honeybees. Journal of Comparative Physiology A, 201(6), 563–584. 10.1007/s00359-015-0987-3.

28. Cashmore, A. R. (2003). Cryptochromes: Enabling plants and animals to determine circadian time. Cell, 114(5), 537–543. 10.1016/S0092-8674(03)00643-4.

29. Ritz, T., Thalau, P., Phillips, J. B., Wiltschko, R., & Wiltschko, W. (2004). Resonance effects indicate a radical-pair mechanism for avian magnetic compass. Nature, 429(6988), 177–180. 10.1038/nature02534.

30. Engels, S., Schneider, N. L., Lefeldt, N., Hein, C. M., Zapka, M., Michalik, A., … & Mouritsen, H. (2014). Anthropogenic electromagnetic noise disrupts magnetic compass orientation in a migratory bird. Nature, 509(7500), 353–356. 10.1038/nature13290.

31. Favre, D. (2011). Mobile phone-induced honeybee worker piping. Apidologie, 42(3), 270–279. 10.1007/s13592-011-0016-x.

32. Helfrich-Förster, C. (2001). The Locomotor Activity Rhythm of Drosophila melanogaster is Controlled by a Dual Oscillator System, Which Is Differently Phase-Shifted by Light and Temperature. Journal of Biological Rhythms, 16(6), 563–573.

33. Tomioka, K., & Chiba, Y. (1982). A Dual Oscillator Model of the Circadian System of the Cricket, Gryllus bimaculatus. Journal of Insect Physiology, 28(12), 997–1006.

34. Beekman, M., & Lew, J. B. (2007). Foraging in honeybees—when does it pay to dance? Behavioral Ecology, 19(2), 255–262.

35. Ingram, K. K., & Gordon, D. M. (2005). Genetic analysis of division of labor in colony-level circadian rhythms in ants. Proceedings of the National Academy of Sciences, 102(21), 7437–7442.

36. Tomioka, K., & Matsumoto, A. (2010). A comparative analysis of circadian organization in insects. Cellular and Molecular Life Sciences, 67(8), 1397–1406.

37. Potts, S. G., Biesmeijer, J. C., Kremen, C., Neumann, P., Schweiger, O., & Kunin, W. E. (2010). Global pollinator declines: Trends, impacts and drivers. Trends in Ecology & Evolution, 25(6), 345–353. 10.1016/j.tree.2010.01.007.

